# Genomics reveals introgression and purging of deleterious mutations in the Arabian leopard (*Panthera pardus nimr*)

**DOI:** 10.1101/2022.11.08.515636

**Authors:** Gabriel Riaño, Claudia Fontsere, Marc de Manuel, Adrián Talavera, Bernat Burriel-Carranza, Héctor Tejero-Cicuéndez, Raed Hamoud M. AlGethami, Mohammed Shobrak, Tomas Marques-Bonet, Salvador Carranza

**Author notes:** Senior authors (Mohammed Shobrak, Tomas Marques-Bonet, Salvador Carranza).

## Abstract

Low genetic variation and high levels of inbreeding are usually a consequence of recent population declines in endangered species. From a conservation point of view, it is essential to genetically screen endangered populations to help assess their vulnerability to extinction and to properly create informed management actions towards their conservation efforts. The leopard, *Panthera pardus*, is a highly generalist predator with currently eight different subspecies inhabiting a wide range of habitats. Yet, genomic data is still lacking for the Critically Endangered Arabian leopard (*P. p. nimr*). Here, we sequenced the whole genome of two specimens of Arabian leopard and assembled the most complete genomic dataset for leopards to date, including genomic data for all current subspecies. Our phylogenomic analyses show that leopards are divided into two deeply divergent clades, one including the only African subspecies and a second one including all seven subspecies of Asian leopards. Interestingly, the Arabian leopard represents a well-differentiated lineage sister to the rest of Asian subspecies. The demographic history, genomic diversity, Runs of Homozygosity (RoHs), and mutational load in the Arabian leopard indicate a prolonged population decline, which has led to an increase in inbreeding and RoHs, with consequent purging of deleterious mutations. Our study represents the first attempt to genetically inform captive breeding programs for this Critically Endangered subspecies. Still, more genomes, particularly from wild individuals, are needed to fully characterise the genetic makeup of this singular and iconic subspecies.

## Introduction

Low genetic variation and high levels of inbreeding are usually a consequence of population decline in endangered species, which may have negative effects on their adaptive potential and rates of reproduction, and thus increase their extinction risk (Frankham et al., 2002). From a conservation point of view, exploring the fine-scale population structure, genetic diversity, and intraspecific demographic dynamics in an endangered species is of crucial importance to correctly design plans for its conservation (Frankham et al., 2002; Shafer et al., 2015). For instance, populations within a species may have different evolutionary histories, substructure or genetic adaptations to their local environment and, if so, they should be considered as different conservation units (Palsbøll et al., 2007). Genetic data plays a key role in detecting all these factors and inferring their effect on demographic changes and inbreeding. Particularly, the use of genome-wide approaches are highly recommended in captive breeding programs, as such datasets can help to identify deleterious mutations and guide the management of endangered species (Irizarry et al., 2016; Johnson et al., 2005; Kohn et al., 2006; Romanov et al., 2006; Shafer et al., 2015; Wright et al., 2020).

In the field of conservation genetics, a few nuclear and mitochondrial markers on a high number of individuals have been used as the standard methodology to infer population parameters (Luikart et al., 2003). These techniques, although very useful, lack the power and precision to reflect all the genomic information from both individuals and populations (Camacho-Sanchez et al., 2020). With the rise and wide implementation of next-generation sequencing (NGS) technologies, the field’s paradigm is shifting towards conservation genomics (Khan et al., 2016). This emerging discipline takes advantage of genome-wide data, such as whole-genome sequencing (WGS), to assess with unprecedented resolution and accuracy both the taxonomy of a focal species, as well as population dynamics such as hybridization events, demographic changes, disease outbreaks and local genetic adaptation (Cho et al., 2013; Figueiró et al., 2017; Fontsere et al., 2022; Frandsen et al., 2020; Kim et al., 2016; Lorenzana et al., 2021; Supple & Shapiro, 2018). More importantly, endangered species are elusive and usually found in low densities, making the task of finding a high number of individuals for multi-locus approaches nearly impossible (Shafer et al., 2015). Sequencing the genome of few individuals has yielded results comparable to those obtained by genotyping a high number of individuals with traditional markers (Irizarry et al., 2016), making WGS a powerful tool for the conservation of endangered species (Wright et al., 2020).

The leopard (*Panthera pardus* Linnaeus 1758) is a highly generalist species, present in a wide range of ecological conditions such as semi-desert, savannah, rainforest and montane habitats, from sea level to high mountain ranges (Miththapala et al., 1996; Uphyrkina et al., 2001). Its diverse diet and capacity to adapt to different environments (Hayward et al., 2006) has allowed this species to expand across Africa and Asia (Miththapala et al., 1996), and even across Europe during the Early Pleistocene (Diedrich, 2013), representing the species with the largest distributional range within the genus *Panthera*. Currently, leopards are still present across much of the African continent and from the Middle East to the Pacific Ocean in Asia (Miththapala et al., 1996; Paijmans et al., 2021; Uphyrkina et al., 2001), although they are only occupying around 25-37% of their historical range (Jacobson et al., 2016). A pioneer study with mitochondrial markers by Uphyrkina et al., (2001) found two main monophyletic groups: the African and the Asian leopards. Fossil evidence and high levels of genetic diversity pointed to an Eastern African origin for the species around 2 million years ago (Mya) (Jacobson et al., 2016; Paijmans et al., 2018; Uphyrkina et al., 2001). However, palaeontological data suggested that more than one out-of-Africa event occurred, and that the Javan leopard (*P. p. melas*) could represent an isolated population resulting from one of those events (Hemmer, 1979; Paijmans et al., 2018; Uphyrkina et al., 2001). Using mitogenomes from historical and ancient samples, a single out-of-Africa dispersion was proposed and dated around 710 thousand years ago (Kya) (Paijmans et al., 2018). Studies with WGS data supported a similar scenario, with a split between African and Asian leopards around 500-600 Kya, and confirmed the monophyly of both groups (Paijmans et al., 2021). Nevertheless, the origin of the Asian colonisation is still uncertain. Paijmans et al., (2021) suggested a colonisation by north-western African leopards. Nonetheless, none of these studies incorporated samples from the Arabian leopard (*P. p. nimr*), a subspecies that due to its geographic distribution is key to resolve the uncertainty on how leopards colonised Eurasia.

The Arabian leopard (*P. p. nimr*) is the Arabian flagship predator and has been listed as Critically Endangered by the IUCN’s Red List of Threatened Species (Mallon & Budd, 2011). This subspecies faces a significant reduction in population size and is on the brink of extinction, with current estimates of fewer than 250 individuals in the wild (Islam et al., 2017; Jackson et al., 2011). Moreover, the Arabian leopard has lost as much as 98% of its historical range (Jacobson et al., 2016), with populations highly isolated and fragmented. This has prevented acquiring knowledge on the current status of wild populations. Previous evaluations estimated 50 individuals in Saudi Arabia, 25-30 individuals in Oman, and a captive stock of 82 individuals, mostly in United Arab Emirates (UAE) (Budd & Leus, 2013; Islam et al., 2017; Mallon & Budd, 2011; Perez et al., 2006). Populations within Yemen have not been evaluated. Furthermore, just nine percent of its current distribution is within protected areas (Jacobson et al., 2016). Nowadays, several wildlife research centres in the Arabian Peninsula are focused on captive breeding programs for this species (Budd & Leus, 2013). However, conservation management efforts for these populations are being formulated without a complete understanding of population genomic patterns, which could result in sub-optimal conservation outcomes (Supple & Shapiro, 2018). Thus, an extensive study of the genomic diversity and population structure of the Arabian leopard and its relationships with other leopard subspecies will strongly benefit its conservation, as a comprehensive knowledge of a species’ history is essential for both evolutionary research and conservation management (Supple & Shapiro, 2018).

Here, we sequence for the first time two whole genomes for the Critically Endangered Arabian leopard at medium coverage (10.31x and 7.52x, respectively) in order to explore its past evolutionary history. Together with the recently released genomes of all currently accepted leopard subspecies (a combined dataset from Paijmans et al., 2021 and Pečnerová et al., 2021), we investigate the phylogenetic position of the Arabian leopard, a highly discussed topic in the past (Paijmans et al., 2021; Uphyrkina et al., 2001). Moreover, we explored introgression between the different subspecies of leopards with special focus between the Arabian leopard with both the African (*P. p. pardus*) and the Anatolian (*P. p. tulliana*) leopards. Finally, with the high resolution provided by WGS data, we assess the current levels of genomic diversity and mutational load, setting the first step for a genomic-informed conservation strategy for the Critically Endangered Arabian leopard.

## Materials and methods

### Data collection

We generated WGS data for two Arabian leopards (*Panthera pardus nimr*): a male and a female from the captive breeding program at Prince Saud Al Faisal Wildlife Research Centre, National Center for Wildlife / the Royal Commission for AlUla (RCU). The male (NIM1 – Arabian1) is the third generation of the captive breeding program whilst the female (NIM2 - Arabian2) is the fourth generation of the same program. Genomic DNA was extracted from whole blood samples using the MagAttract HMW Kit (Qiagen) following manufacturer’s protocols. Then, we prepared Illumina libraries following the BEST protocol (Carøe et al., 2018) with minor modifications. The libraries were sequenced in a 2×101 bp HiSeq4000 lane aiming for a 10x depth of coverage. Additionally, we gathered the raw sequencing data (FASTQs) from 18 leopards of all the other subspecies and three lions (*Panthera leo*) (See Table 1) generated in previous publications (de Manuel et al., 2020; Kim et al., 2016; Paijmans et al., 2021; Pečnerová et al., 2021). Additionally, we obtained two full mitochondrial genomes for two extinct Pleistocene European samples (*P. p. spelaea*) from Paijmans et al., (2018).

**Table 1:**
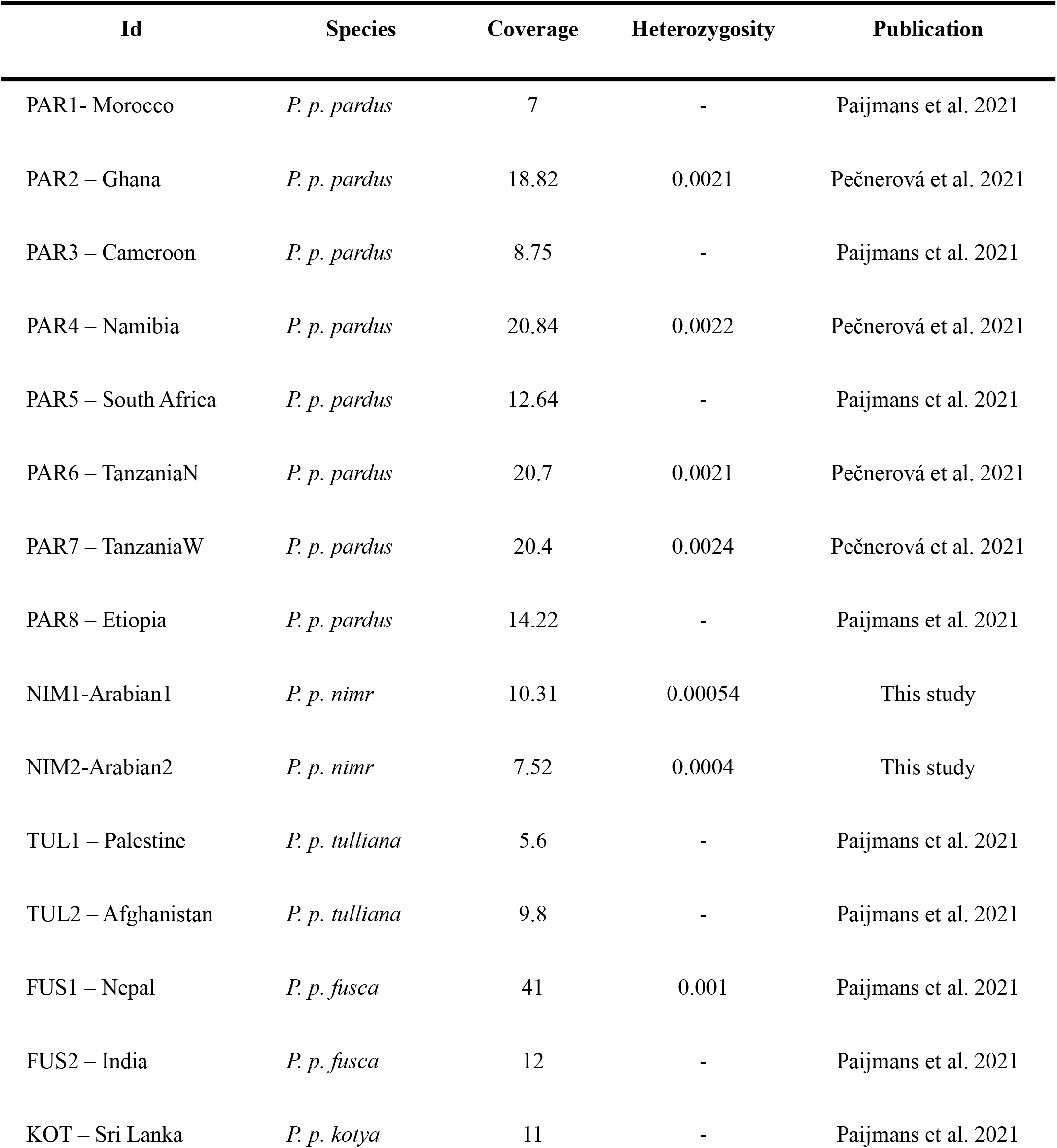
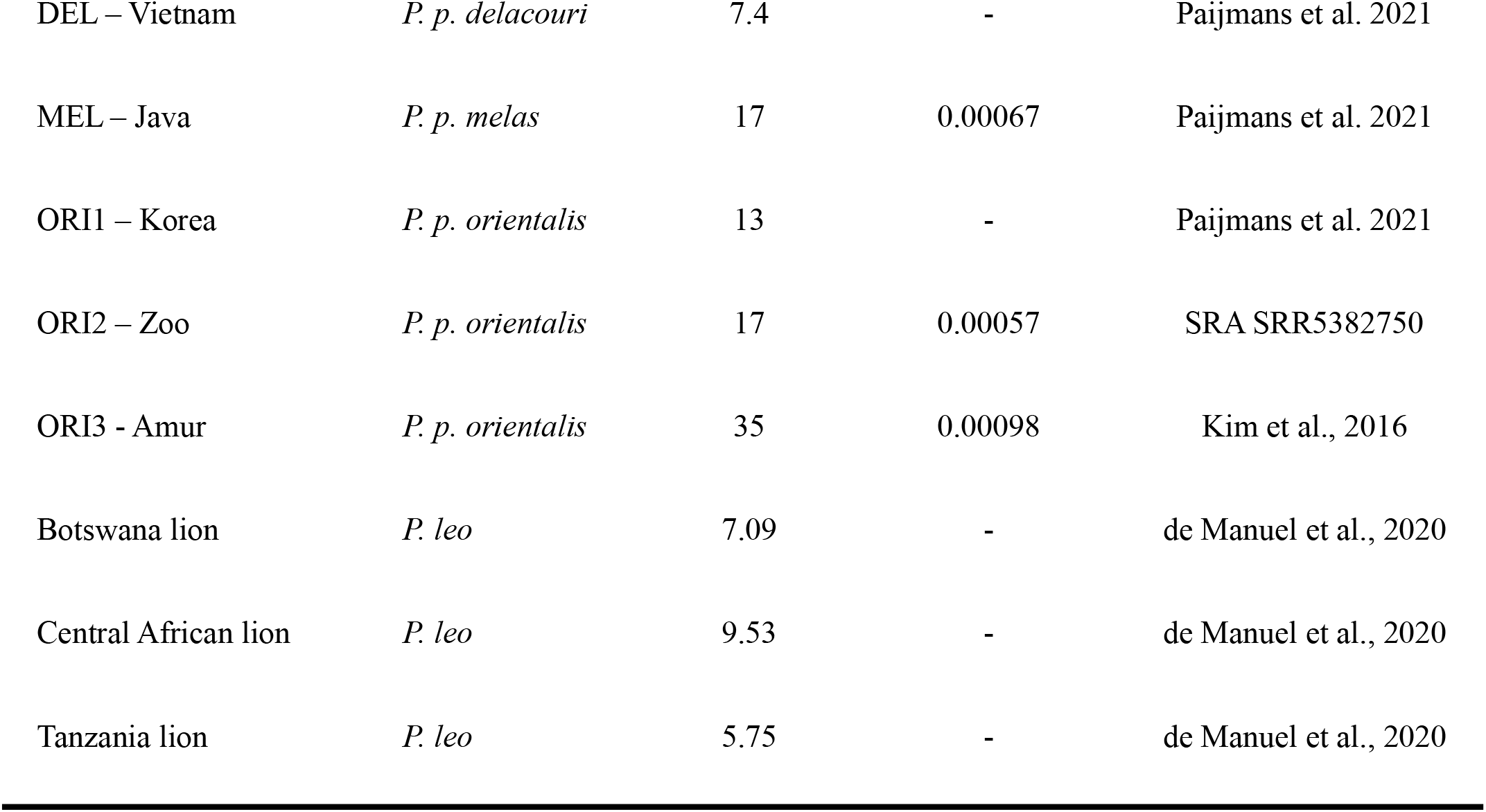
Information of all samples used in this study, including coverage and heterozygosity for species with coverage higher than 15x.

### Data processing

Raw reads for each of the 23 genomes were filtered, and adapters removed with fastp (Chen et al., 2018). A minimum base quality score was set to 30, and adapter detection for paired-end sequencing was activated, with a required fragment length of 50bp. Trimming of poly-G/X tails and correction in overlapped regions were specified. All other parameters were set as default. Filtered sequences were visually explored with FastQC (Andrews, 2010) to ensure data quality and absence of adapters. Filtered reads were mapped against the reference genome of a female domestic cat (Felis_catus_9.0; GenBank assembly accession: GCA_000181335.4) (Buckley et al., 2020) with bwa-mem v0.7.17 (H. Li, 2013). Mapped reads were sorted with Samtools v1.9 (Li et al., 2009). Duplicated reads were marked and removed with PicardTools (Broad Institute, 2021) and reads with mapping quality lower than 30 were discarded. SNP calling was carried out with HaplotypeCaller from GATK (McKenna et al., 2010), with BP_resolution and split by chromosome. For each chromosome, individual genotypes were joined using CombineGVCFs with convert-to-base-pair-resolution and the GenotypeGVCFs tool was then applied to include non-variant sites. Finally, the whole dataset split by chromosome was concatenated with bcftools concat (Danecek et al., 2021), keeping only the autosomes. For some analyses we generated a separate dataset including only leopards. Then, we filtered the raw callset by excluding variants matching at least one of the following criteria: Quality by Depth (QD) <10, Mapping Quality (MQ) < 50, Fisher Strand test (FS) > 10, StrandOddsRatio (SOR) > 4, MQRankSum < −5 && MQRankSum > 5 and ReadPosRankSum < −5 && ReadPosRankSum > 5. Later, genotypes were filtered using vcftools (Danecek et al., 2011), with a minimum variant quality of 30, removing indels, keeping only biallelic SNPs, allowing 10% missing data and filtering for minor allele frequency (MAF) of 0.001, filtering out monomorphic sites and keeping only variable positions. Repetitive regions were identified from the cat reference genome and removed. A second dataset was created for population genetic analyses keeping only unlinked SNPs through bcftools (Danecek et al., 2021) using a maximum value of r^2^= 0.5.

### Population structure analyses

We performed a Principal Component Analysis (PCA) with the leopard unlinked filtered dataset using Plink v1.9 (Chang et al., 2015). We applied ADMIXTURE (Alexander et al., 2009) to detect ancestral populations from k=2 to k=5. A total of 20 replicates for each K were calculated, selecting the best K value after 20 cross validations. Visualisation of results from these analyses was performed with R v.3.6.3 (R Core Team, 2021) and the R package ggplot2 (Wickham, 2016).

### Phylogeny

We reconstructed the phylogenomic relationships of all samples (including the three lion genomes) using only autosomal chromosomes selected from bam files with the view function from Samtools v1.9 (Li et al., 2009). These bam files were used to generate individual pseudohaploid consensus sequences with ANGSD v.0.933 (Korneliussen et al., 2014), taking a consensus-based sampling approach (-doFasta 3) in non-overlapping sliding windows of 1 Mbp. Maximum likelihood trees for each window along the cat reference genome were calculated with IQ-TREE2 (Nguyen et al., 2015) applying a GTR+I+G substitution model, with the lion samples as outgroup. Windows where one individual had >50% missing data were removed, leaving a total of 2,330 non-overlapping windows. A maximum clade-credibility tree was created with TreeAnnotator v2.6.4. from BEAST2 v2.6.6 (Bouckaert et al., 2019). We tested the effect of window size by repeating the analyses for smaller window sizes (500Kbp and 100Kbp) for the largest chromosome (240 Mbp) and similar topologies were obtained. Additionally, the mitochondrial DNA (mtDNA) was assembled with MitoFinder v.1.4.1 (Allio et al., 2020) from a subset of the samples and a maximum likelihood phylogeny was reconstructed using RAxML-NG (Kozlov et al., 2019), with the GTR+I+G model and performing 1,000 bootstraps.

### Demographic history

We inferred the demographic history of leopards with the Pairwise Sequential Markovian Coalescent (PSMC) software (Li & Durbin, 2011), for which we used four high-coverage individuals together with the two Arabian leopards. Heterozygous positions were obtained from bam files with Samtools v1.9 (Li et al., 2009) and data was filtered for low mapping (<30) and base quality (<30). Minimum and maximum depths were set at half and double the average coverage for each sample, respectively. Only autosomal data was considered. A rate of 1.1 x 10^-8^ substitutions/site/generation and a generation time of 5 years were used, following (Pečnerová et al., 2021). Other parameters were set as default following previous knowledge on leopard genomics (Kim et al., 2016). For the two Arabian leopards, 100 bootstraps were calculated. Final results were plotted with the psmc_plot.pl function from PSMC (https://github.com/lh3/psmc). Finally, high-coverage individuals were downsampled to the coverage level of the Arabian leopards and PSMC was run again, with comparative purposes.

### Genomic diversity and RoHs

We used the raw dataset to calculate average genome heterozygosity per individual. We generated non-overlapping sliding windows of 100 Kbp for the domestic cat reference genome and took only sites (both variant and invariant) with site quality higher than 30 (QUAL field in a VCF file from GATK). Only windows containing more than 60,000 unfiltered sites were considered. Later, six individuals were downsampled to coverage levels similar to those of the Arabian leopards, for comparative reasons. After obtaining homozygous regions per individual (see below), out-of-RoH heterozygosity was calculated using the intersect function from bedtools (Quinlan & Hall, 2010). Runs of Homozygosity (RoHs) were calculated per individual based on the density of heterozygous sites in the genome using the implemented Hidden Markov Model (HMM) in bcftools roh function (Narasimhan et al., 2016) with –AF-dflt 0.4 (following Armstrong et al., 2020) on the filtered dataset. We kept RoHs with a Phred Score of at least 70 and with a minimum length of 100 Kbp. Finally, we performed a correlation test between the number and cumulative length of RoHs. Visualisation for all analyses was carried out with ggplot2 (Wickham, 2016).

### Introgression

We used the D-Statistics (ABBA-BABA tests) method using the qpDstat function from Admixtools (Patterson et al., 2012) to test for gene flow between the different subspecies of leopards and especially between the Arabian leopard and its geographically closest subspecies of leopards (i.e., African and Anatolian leopards). To do so, we used the filtered VCF file (16.32 million SNPs) containing all subspecies of leopards and using the lion as an outgroup. We first explored the possible introgression between African and Arabian leopards, as previous mitochondrial phylogenetic analyses showed African and Arabian leopards clustering together (Uphyrkina et al., 2001). When introgression between African and Arabian leopards was tested, we created a model with (((X, Arabian),African),Lion), where X are all other subspecies of leopards. Posterior comparisons used the same procedure, changing the order of the samples accordingly. Later, and following population genetic analyses, we examined if the Arabian leopards contained past signals of introgression with the Anatolian leopard. Finally, we tested the subsequent possible introgressions between the remaining subspecies.

### Mutational load

We estimated the mutational load for coding regions in all leopard individuals using SNPeff v.4.3. (Cingolani et al., 2012) with the filtered dataset, not allowing missingness (10.35 million SNPs). A database was created using the cat annotation file available in GenBank (Felis_catus_9.0; GenBank assembly accession: GCA_000181335.4) (Buckley et al., 2020). We identified putative deleterious variants in the four categories established by the manual: 1) Low: mostly harmless or unlikely to change protein behaviour (i.e., synonymous variants); 2) Moderate: non-disruptive variants that might change protein effectiveness (i.e., missense variants); 3) High: assumed to have a high (disruptive) impact in the protein, probably causing protein truncation, loss of function (LoF) or triggering nonsense mediated decay (i.e., stop codons, splice donor variant and splice acceptor, start codon lost, etc.); 4) Modifier: usually non-coding variants or variants affecting non-coding genes, where predictions are difficult or there is no evidence of impact (i.e., downstream or upstream variants) (Cingolani et al., 2012). Then, following the author’s recommendations (Cingolani et al., 2012), we counted the number of derived alleles based on the cat reference genome with low, moderate and high predicted levels for homozygous (multiplied by two as they are represented twice) and heterozygous alleles, removing observations with warnings. Later, we calculated the amount of high-impact deleterious alleles in and outside of RoH regions using bcftools. Because we only used sites found in all individuals, variants in these two categories were separately counted and no bootstrapping was needed, as discussed in Dussex et al., (2021). Following Xue et al., (2015), we calculated a statistic which compares two populations in relation to the number of derived alleles found at sites in one population rather than the other, for a particular variant category. Basically, for each category of variants we estimated at each site *i* the observed allele frequency in Population X as f^x^_i_= d^x^_i_ / n^x^_i_, where n^x^_i_ is the total number of alleles called in population X and d^x^_i_ is the total number of called derived alleles. Similarly, we define f^y^_i_ for population Y. Then, for each category C of variants, we estimated:

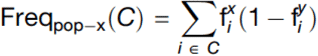

Finally, we calculated the ratio R_XY_ = Freq_pop-x_ / Freq_pop-y_ where a value of 1 corresponds to no change in frequency, R_XY_ > 1 represents a decrease in frequency in population Y relative to population X and R_XY_ < 1 results from an increase in frequency in population Y relative to population X.

## Results

### Population structure

We generated the whole genome from two Critically Endangered Arabian leopards at an average coverage of 10.32x and 7.53x (Table 1). Together with the African and Asian samples gathered from previous studies (Kim et al., 2016; Paijmans et al., 2021; Pečnerová et al., 2021), we assembled a genomic dataset for the leopard lineage that, for the first time, includes representatives from all current subspecies (Kitchener et al., 2017) (Fig. 1A and Table 1). With this comprehensive dataset, we explored the population structure of leopards with a Principal Component Analysis (PCA) of 1.35 million polymorphic positions, after LD pruning. We found a segregation of African and Asian populations along PC1 (Fig. 1B). PC2 informed about the variability within the Asian specimens, recovering a gradient following geographic locations. Interestingly, the Arabian leopards clustered next to their geographically closest populations from Palestine and Afghanistan. Similarly, Admixture analyses reported k=2 as the most likely number of ancestral populations (Table S1), dividing African and Asian populations (Fig. 1C). Arabian leopards grouped within the Asian cluster for k=2 however showing up to 37% of African component. The Arabian, the Palestinian and the Afghanistan leopards formed a unique cluster for k=3 (the latter with 41% of ancestral component from the Asian clade).

**Fig. 1:**
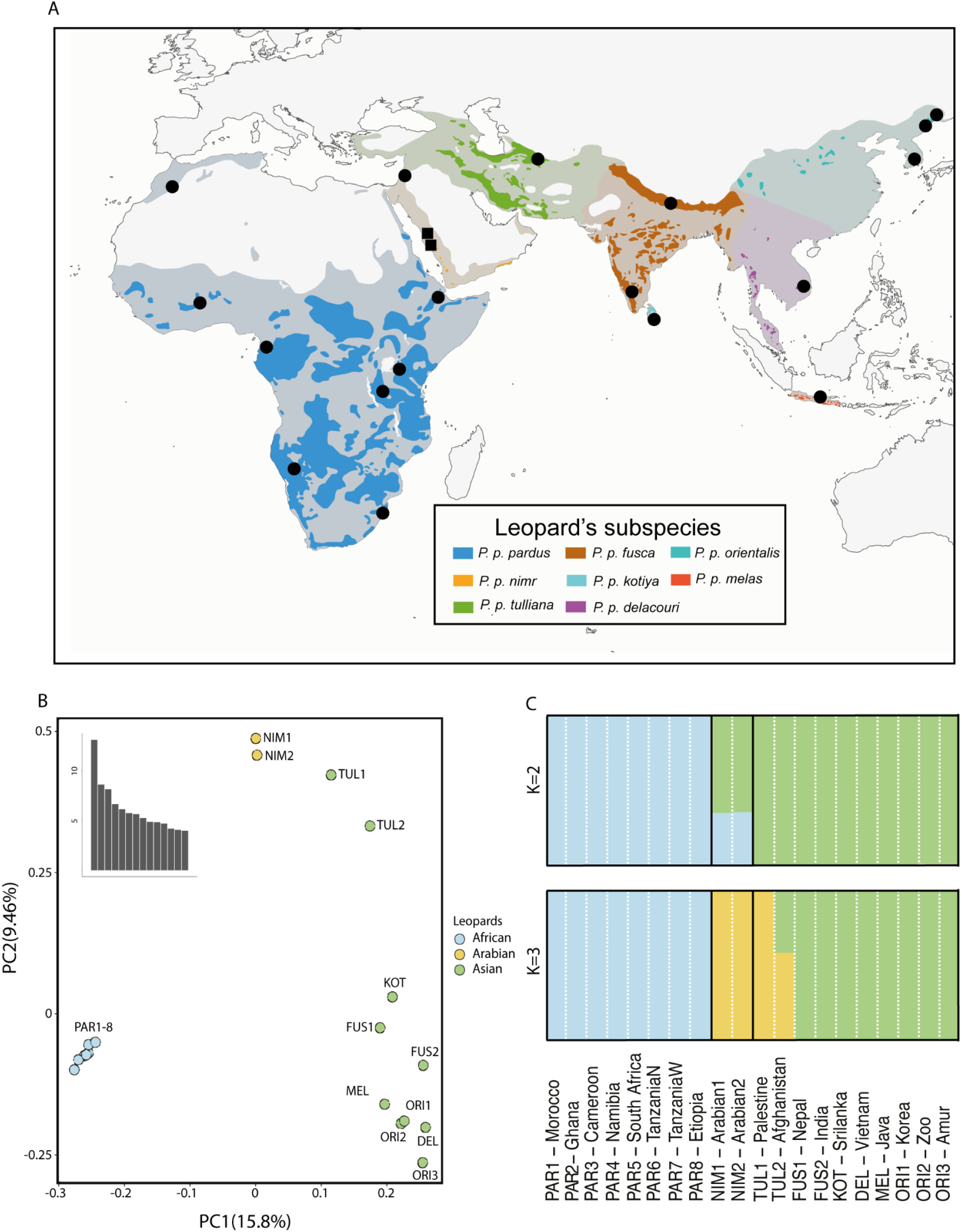
Sampling and population genetic structure. (A) Distribution for all non-extinct leopard subspecies, including historical distribution for all the species (in lighter colour). Black squares show new individuals sequenced for this study whilst circles show data already available and included in this work. (B) PCA of genetic variation over 1.35 million SNPs for all subspecies of leopards. (C) Admixture analysis with k=2 and k=3. Abbreviations are as follows: PAR, *P. p. pardus*; NIM, *P. p. nimr*; TUL, *P. p. tulliana*; FUS, *P. p. fusca*; KOT, *P. p. kotiya*; DEL, *P. p. delacouri*; ORI, *P. p. orientalis* and MEL, *P. p. melas*.

### Evolutionary relationships

To explore in more detail the evolutionary relationships of all leopard subspecies and in particular to shed light on the phylogenetic position of the Arabian leopards, we inferred their phylogenetic relationships using 2,330 non-overlapping 1 Mbp windows that covered around 96% of the whole genome. The resulting phylogenomic tree shows two well-supported main clades, the African and the Asian (Fig. 2A). Although the internal phylogenomic relationships within the African clade were not well supported (a result previously discussed by Paijmans et al., 2021), the internal phylogenomic relationships within the Asian clade support the Arabian leopard as sister taxon to all the remaining subspecies of Asian leopards. All the other phylogenomic relationships within the Asian clade were not fully supported, and the analysis did not recover monophyly for some of the subspecies of Asian leopards (Fig. 2A). Phylogenomic trees with sliding windows of reduced size (i.e. 500 Kbp and 100 Kbp) showed similar topologies with overall lower support (although some subspecies within the Asian clade were not recovered as monophyletic) (Fig. S1). In both cases, Arabian leopards were found within the Asian clade. Interestingly, the phylogenomic tree with a sliding window of 100 Kbp showed one individual of the Anatolian leopard (*P. p. tulliana*) clustering together with the Arabian leopards, a result in line with introgression analyses (see below). The mitogenomic phylogeny split the samples into two main groups: a well-supported clade comprising the African and Arabian subspecies (contrary to the nuclear genome phylogeny of Fig. 2A), and an Eurasian clade comprising European samples (*P. p. spelaea*) from the late Pleistocene and all other remaining leopard Asian subspecies, showing a topology similar to previous phylogenies (Paijmans et al., 2018; Uphyrkina et al., 2001) (Fig. 2B).

**Fig. 2:**
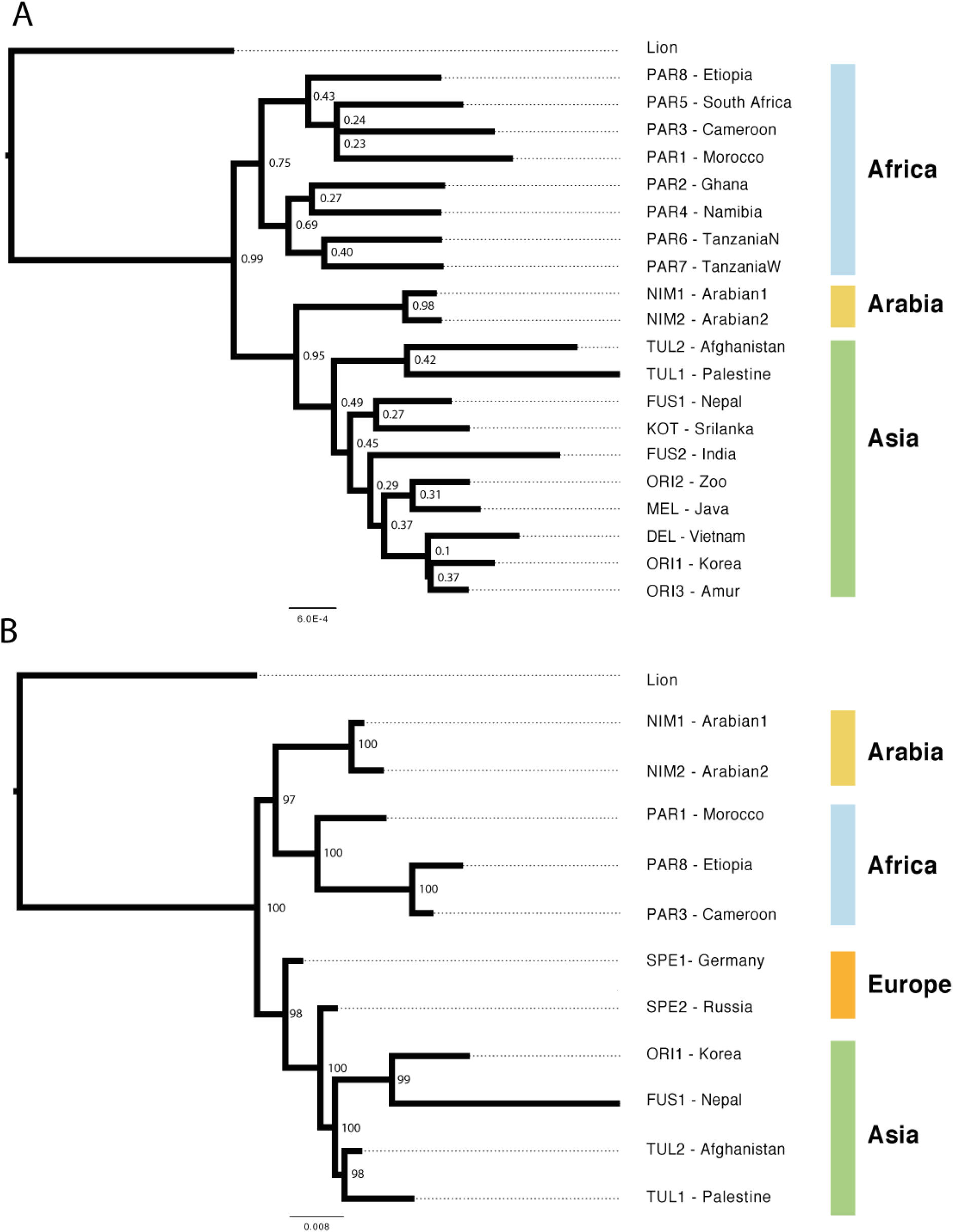
Phylogenomic trees for the nuclear and mitochondrial data. (A) Phylogenomic consensus tree from 2,330 Maximum Likelihood trees of 1 Mbp non-overlapping sliding windows along the reference genome for all currently accepted subspecies of leopards, and with the lion as outgroup. Node values indicate clade frequency. (B) Maximum likelihood mitogenome phylogeny for a subset of the samples, with lion as outgroup. Bootstrap values are shown in each node. Abbreviations are as follows: PAR, *P. p. pardus*; NIM, *P. p. nimr*; TUL, *P. p. tulliana*; FUS, *P. p.fusca*; KOT, *P. p. kotiya*; DEL, *P. p. delacouri*; ORI, *P. p. orientalis*; MEL, *P. p. melas* and SPE, *P. p. spelaea*.

### Ancient demographic history

PSMC analysis showed a continuous negative trend in population size for the two Arabian leopards (Fig. 3 and Fig. S2). Individuals from other subspecies showed results previously observed (Paijmans et al., 2021; Pečnerová et al., 2021), with African samples reporting the highest current effective population size of the species and Javan and Amur samples showing lower population sizes, with the Amur leopard following an ancestral trajectory different from African samples, as reported in Paijmans et al., (2021). Due to the lower coverage of both Arabian leopards compared to the other subspecies, we downsampled the high-coverage data at similar coverage levels to those of the Arabian leopards (Fig. S3). As expected, lower effective population sizes were observed with the downsampled dataset but general trends persisted (compare Fig. 3 with Fig. S3), indicating that the pattern observed in the Arabian samples was likely not strongly affected by their coverage levels. Beyond >10^5^ years, PSMC inferred different effective population sizes for the two Arabian leopards, most likely the product of inaccuracy due to their low heterozygosity and lack of coalescent events dating at that point in the past (Li & Durbin, 2011).

**Fig. 3:**
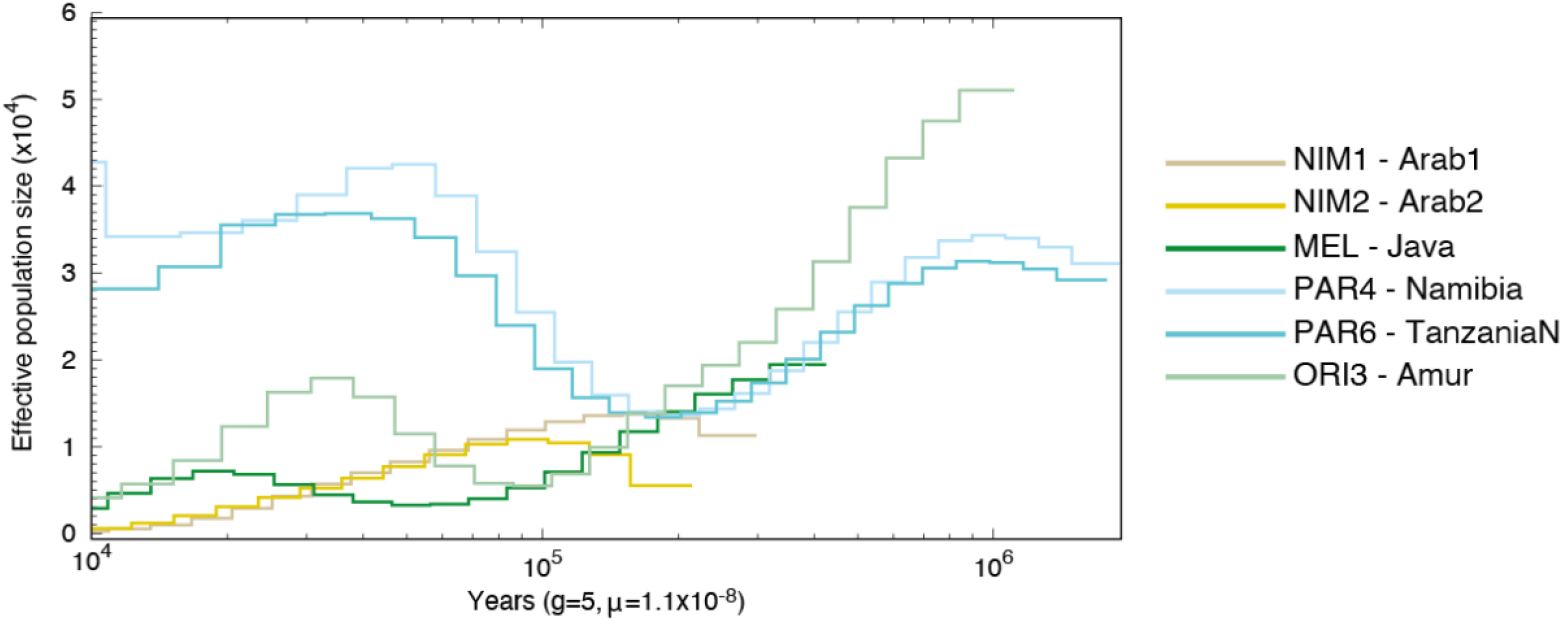
PSMC analysis for the high-coverage samples plus the two Arabian leopards. Generation time was set to 5 years and substitution rate to 1×10^-8^ per site per year. Abbreviations are as follows: NIM, *P. p. nimr*; MEL, *P. p. melas*; PAR, *P. p. pardus* and ORI, *P. p. orientalis*.

### Genomic diversity, inbreeding and RoHs

Genome-wide heterozygosity levels varied across subspecies, with the African lineage harbouring the highest diversity levels (Fig. 4A). Heterozygosity in Asian individuals varied among the subspecies, with the Arabian leopards having some of the lowest heterozygosity values (5.4×10^-4^ and 4×10^-4^ for Arabian1 and Arabian2, respectively). Individuals within the Amur leopard subspecies (*P. p. orientalis*), classified as Critically Endangered by IUCN, showed in general low heterozygosity but higher than the Arabian samples, whilst the Nepalese leopard (*P. p. fusca*) contained the highest heterozygosity values within the Asian clade. Out-of-RoH heterozygosity recovered higher estimates but showed similar tendencies to genome-wide heterozygosity, with Arabian leopards showing more than double their genome-wide heterozygosity (Fig. S4). To test the effect of coverage on the detection of heterozygous sites, we downsampled six high-coverage individuals to similar coverage levels to the Arabian leopards and we did not detect any statistically significant change between non-downsampled and downsampled individuals (t = −0.66, *p*-value = 0.51), although the resulting heterozygosity levels were slightly lower after downsampling, as expected (Fig. 4A). After correcting for the coverage, Arabian leopards still showed very low heterozygosity levels, with significant differences with downsampled African (t = 16.61, *p*-value = 0.007) but not with other downsampled Asian leopards (t = 0.62, *p*-value = 0.57). RoHs analyses reported high disparity among all the samples tested (Fig. 4B). Both Arabian leopards contained more than 50% of the genome under RoH (Fig. 4B; Table S2), with the highest percentage of short (<500 Kbp) and medium (0.5-1 Mbp) RoHs, indicative of a small long-term effective population size. Moreover, there was an important difference in the number of long RoHs (>1 Mbp) between the two Arabian leopards (Fig. 4B; Table S2). This could be explained by different levels of recent inbreeding as both individuals are captive-bred leopards. Arabian2, a fourth-generation captive-bred female, had 10% longer RoHs than Arabian1, a third-generation captive-bred male. The Javan leopard had around 30% of short RoHs, indicating ancient inbreeding in this island subspecies. Interestingly, one captive *P. p. orientalis* individual (ORI2-Zoo) accumulated almost 50% of its genome within RoHs and 20% of them being long (>1 Mbp). All these results were supported by a correlation between Number (NROH) and cumulative sum of RoHs (SROH) (Pearson correlation t = 3.14, correlation value = 0.76 and *p*-value = 0.016; Fig. S5; Table S3), with large and well connected populations having a reduced NROH and SROH and small and fragmented populations showing high values of both NROH and SROH.

**Fig. 4:**
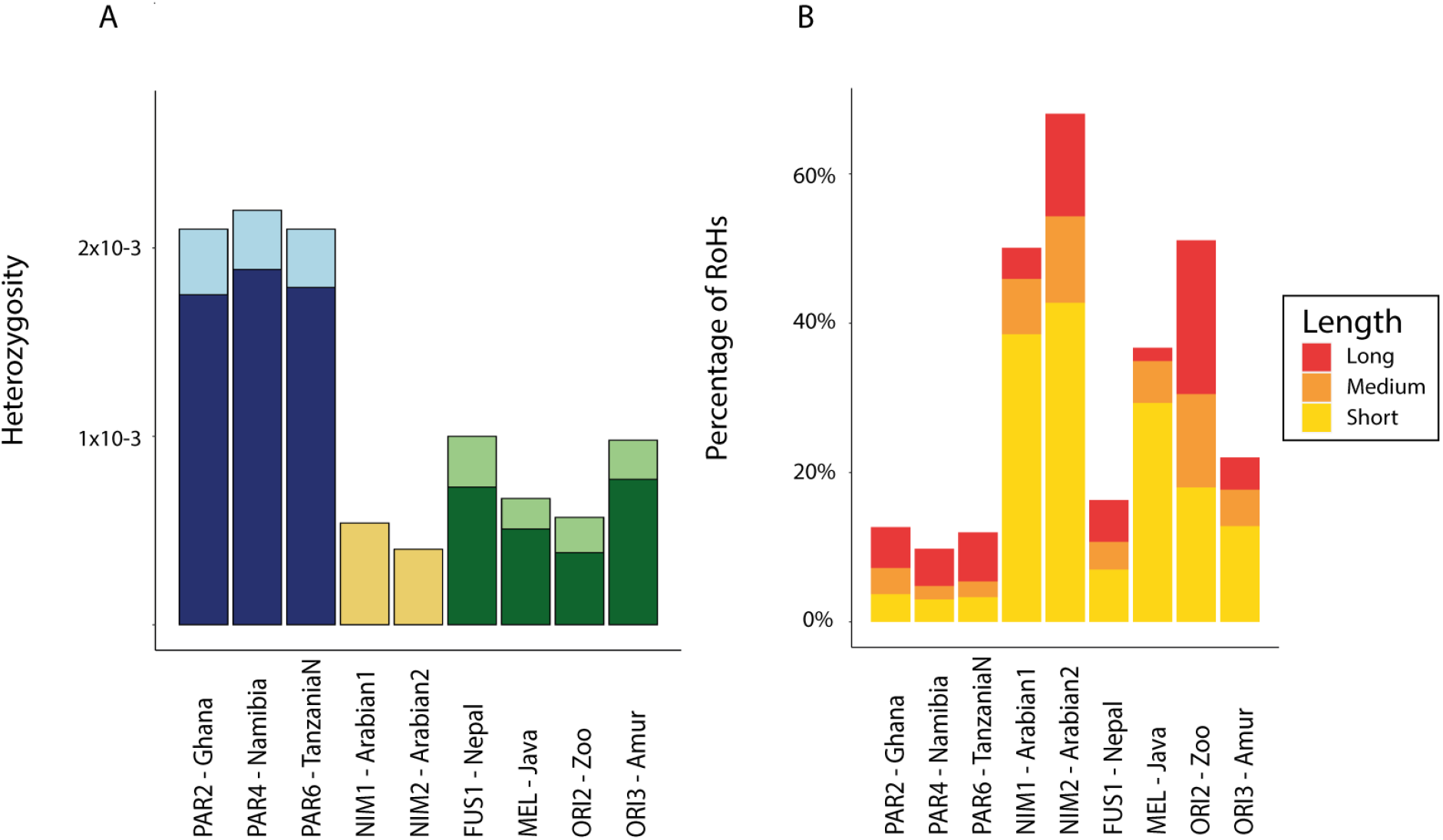
Heterozygosity and Runs of Homozygosity for the high-coverage individuals plus the Arabian leopards. (A) Genome-wide heterozygosity for all samples with coverage levels higher than 15x and the two Arabian leopards. In dark, genome-wide heterozygosity for downsampled individuals, in light heterozygosity with non-downsampled individuals. Arabian leopards were not downsampled. Statistically significant differences exist between downsampled African and Arabian leopards (*p*-value < 0.01) but not between downsampled Asian and Arabian (*p*-value = 0.57). (B) Percentage of the genome in RoH. Different colours within the columns indicate the relative percentage of long (>1 Mbp), medium (>500 Kbp and <1 Mbp) and short (<500 Kbp) RoHs along the genome for the high-coverage samples and the two Arabian leopards; see Table S2 for more detailed information. Abbreviations are as follows: PAR, *P. p. pardus*; NIM, *P. p. nimr*; FUS, *P. p. fusca*; MEL, *P. p. melas* and ORI, *P. p. orientalis*.

### Introgression

We initially tested introgression between the Arabian leopard and its two geographically closest subspecies, the African (*P. p. pardus*) and the Anatolian (*P. p. tulliana*) leopards, and later between all other leopard subspecies. We tested a simple tree-like model where the lion was set as an outgroup following the new phylogenomic relationships to establish the comparisons. Introgression analyses revealed past introgression between Arabian and both African and Anatolian leopards (Fig. 5). D-statistic values varied depending on the comparison, but all of them were significant and with absolute Z-scores higher than three. We also found introgression between several other subspecies, highlighting a complex history of gene flow within the species (Table S4).

**Fig. 5:**
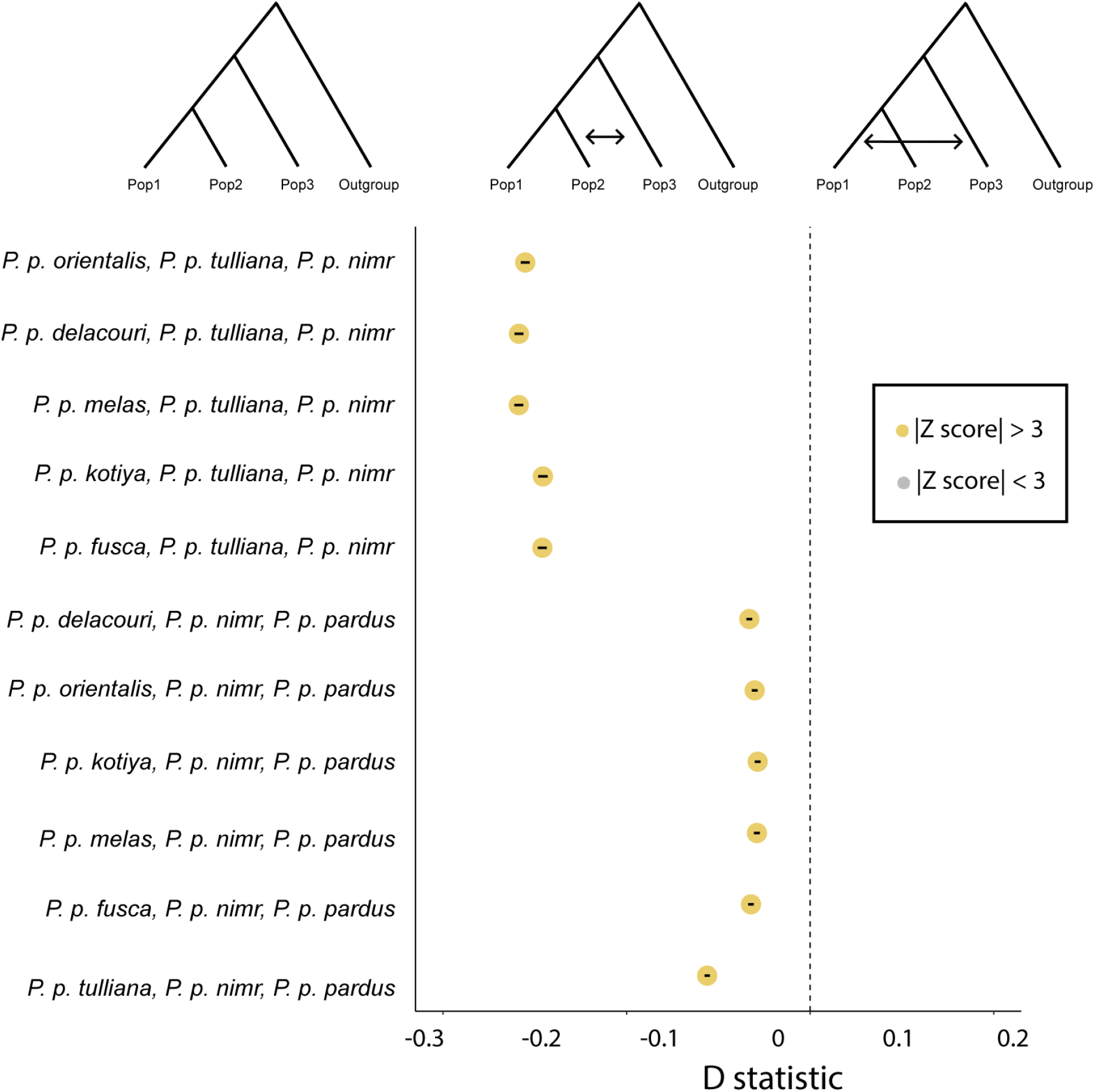
Introgression analyses between the Arabian leopard (*P. p. nimr*) and the two geographically closest subspecies, the African (*P. p. pardus*) and the Anatolian (*P. p. tulliana*) leopards, using the lion as outgroup. Introgression between Pop2 and Pop3 will be reported when D-statistic is negative and introgression between Pop1 and Pop3 will happen when the D-statistic is positive. All D-statistics had an absolute Z-score value higher than 3 (shown in yellow). Error bars are shown within the dots.

### Mutational load

We compared the mutational load of the Arabian leopard (an example of a small and long-term isolated subspecies) to that of the African (a subspecies with large and connected populations) and to other Asian leopards (an intermediate case, with some subspecies with well connected populations and others with small and isolated populations). As a first approach, we counted the number of high, moderate or low impact deleterious alleles for each individual (see Methods). Later, we calculated the number of homozygous (multiplied by two as they are represented twice) and heterozygous alleles separately, as a proxy of realised (i.e., homozygous alleles with effects on the current generation) and masked (heterozygous alleles that can be expressed in future generations) genetic load, thus assuming most deleterious alleles are recessive (Bertorelle et al., 2022). Moreover, we calculated the ratio of derived alleles between African, Arabian and Asian individuals (although Arabian leopards are within Asia, we analysed them separately for comparative reasons). Arabian leopards showed the lowest number of high-impact deleterious alleles but similar levels of moderate and low-impact deleterious alleles compared to African and the rest of Asian leopards (Fig. 6A and Fig. S6). Interestingly, Arabian leopards significantly differed from both African and the rest of Asian leopards in the number of high-impact deleterious alleles (t = 2.93, *p*-value = 0.021 and t = 4.35, *p*-value = 0.002, respectively) and with Asian but not African leopards in the number of moderate-impact deleterious alleles (t = 3.75, *p*-value = 0.005 and t = −0.28, *p*-value = 0.78, respectively) (Fig. 6A and Fig. S6). Interestingly, low-impact deleterious alleles did not show differences between Arabian and African leopards (t = 1.68, *p*-value = 0.13), Arabian and the rest of Asian leopards (t = 1.74, *p*-value = 0.11) and African and Asian leopards (excluding Arabian leopards) (t = 0.39, *p*-value = 0.7). African and Asian (excluding Arabian) leopards did not differ in the number of high-impact deleterious alleles (t = −1.95, *p*-value = 0.07) but did in the number of moderate-impact deleterious alleles (t = −2.66, *p*-value = 0.01). The number of high-impact and moderate-impact derived deleterious alleles in homozygosity (the realised load) showed significantly higher number of alleles for Arabian leopards compared with African leopards (t = −14.24, *p*-value < 0.0001 for high-impact deleterious alleles; t = −26.15, *p*-value < 0.0001 for moderate-impact deleterious alleles) and only for moderate-impact deleterious alleles when compared with other Asian leopards (t = 0.53, *p*-value = 0.60 for high-impact deleterious alleles; t = −2.68, *p*-value = 0.02 for moderate-impact deleterious alleles) (Fig. 6A and Fig. S6). Conversely, Arabian leopards showed significantly lower numbers of heterozygous alleles (the masked load) compared to African (t = 10.7, *p*-value < 0.0001 for high-impact deleterious alleles and t = 13.48, *p* > 0.0001 for moderate-impact deleterious alleles) and the rest of Asian leopards (t = 3.39 *p*-value = 0.09 for high-impact deleterious alleles and t = 3.99, *p*-value = 0.003 for moderate-impact deleterious alleles) (Fig. 6A and Fig. S6). Ratio of derived alleles (R_XY_) between Arabian leopards and both African and the rest of Asian leopards were in line with previous results and showed reduced high-impact and moderate-impact deleterious alleles for the former whilst the ratio between Asian (excluding Arabian) and African leopards showed Asian leopards to have slightly more deleterious alleles for both categories (Fig. 6B). Around half of the observed alleles were within RoHs for the Arabian leopards and the *P. p. orientalis* specimen analysed from the Zoo, whilst all other leopards had at least 75% of the deleterious alleles outside RoHs (Fig. S7).

**Fig. 6:**
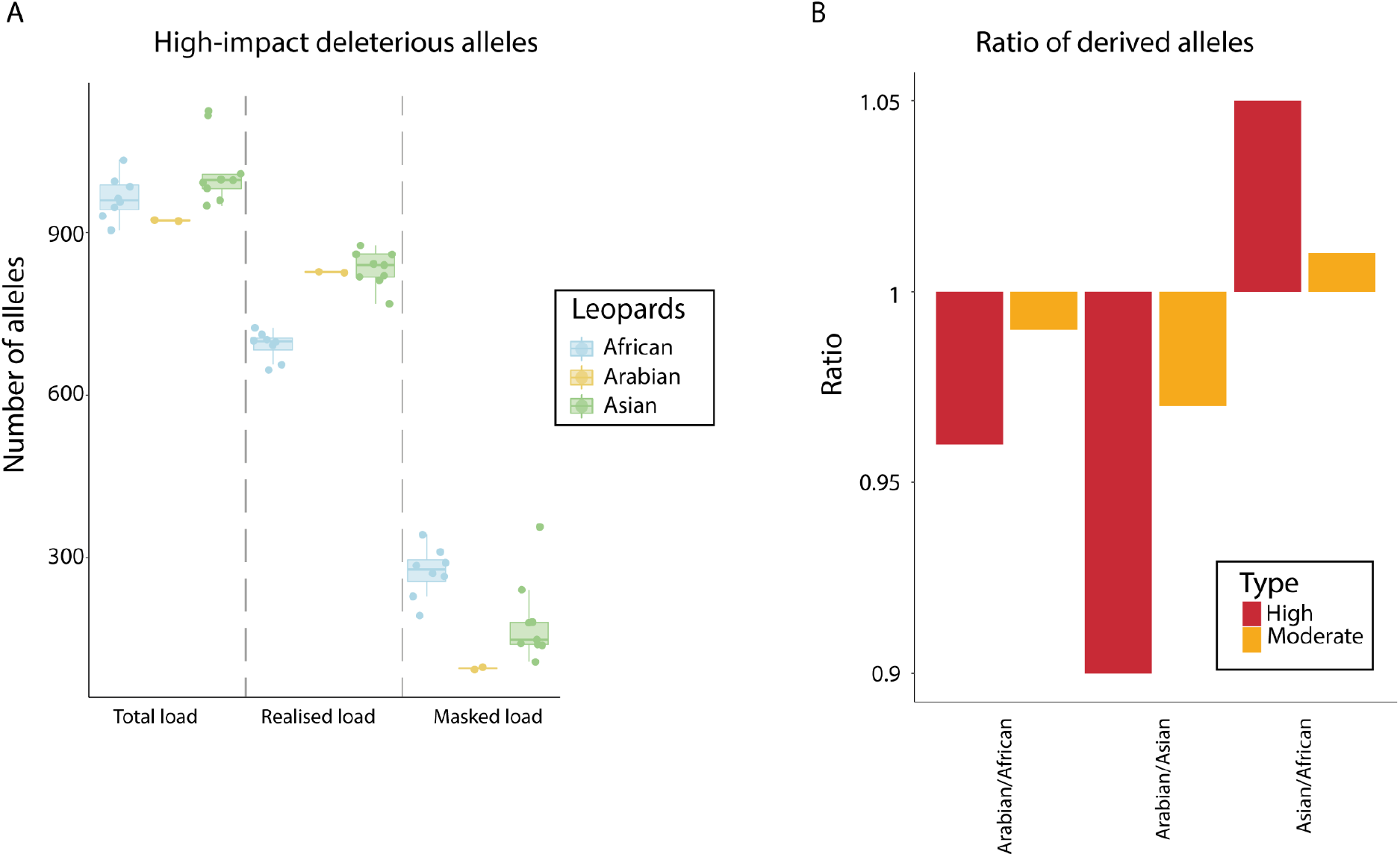
Mutational load in leopards. (A) Number of high-impact deleterious alleles found in total, realised and masked genetic load for leopards from Africa, Arabia and the rest of Asia. (B) Ratio of high and moderate impact deleterious alleles between Arabian leopards with both African and the rest of Asian leopards and between African and Asian leopards (excluding Arabian) for both high and moderate-impact deleterious alleles.

## Discussion

In this study, we have sequenced for the first time the complete genomes of two Arabian leopards (*P. p. nimr*), a Critically Endangered subspecies with less than 250 wild individuals distributed into continuously declining and severely fragmented populations across Saudi Arabia, Yemen and south Oman (Islam et al., 2017; Jackson et al., 2011). Apart from its interest from a conservation point of view, its geographical distribution at the intersection between the two main leopard clades (Africa and Asia) make this subspecies key to understanding the evolutionary history of leopards. Together with genomic data available from all other leopard subspecies (Fig. 1A), we have analysed their evolutionary history and demography. In an attempt to guide future conservation plans in the Arabian leopard and other subspecies, we have applied a conservation genomics approach to assess the genetic consequences of isolation and inbreeding.

Population structure analyses suggest that the Arabian leopards are closer to the Asian than to the African leopards (Fig. 1B and 1C). We replicated the PCA results from Paijmans et al., (2021), with all African leopards clustering very closely both in PC1 and PC2, while the Asian leopard subspecies appeared separated in PC2 according to their geographical distribution. These contrasting clusterings highlight a lower genetic differentiation within African leopards than within Asian leopards (Fig. 1B). Genetic similarity in African leopards can be explained by high mobility, habitat versatility and weak signatures of dispersal barriers across Africa (Jacobson et al., 2016; Pečnerová et al., 2021; Ray-Brambach et al., 2018). Conversely, Asian leopards are characterised by high levels of structuring (Fig. 1B). This is considered as an intrinsic feature of these populations, possibly caused by drift after dispersal into Asia, as expansions can produce population structure at neutral loci (Excoffier & Ray, 2008; Paijmans et al., 2021).

Due to the almost complete absence of genetic information for the Arabian leopard to date, its phylogenetic position has been a highly discussed topic for decades (Paijmans et al., 2021; Pečnerová et al., 2021; Uphyrkina et al., 2001). Previous to this study, the only genetic information available was from a study based on mitochondrial markers suggesting that the Arabian leopard was the sister group to the African leopard (Uphyrkina et al., 2001). Our phylogenomic analysis, including thousands of genome-wide autosomal markers of all currently accepted subspecies of leopards, was key to successfully resolve the evolutionary relationships of leopards and the enigmatic phylogenetic position of the Arabian leopard (Fig. 2A). Contrary to the results of the analysis of the mitochondrial dataset by Uphyrkina et al., (2001), and the results of our mitogenome analyses presented in this study (Fig. 2B), the genome-wide autosomal dataset fully supported the position of the Arabian leopard as sister taxon to the rest of Asian leopards (Fig. 2A); a result consistent with the population structure analyses (Fig. 1), and suggesting an only out-of-Africa dispersal event. This is not the first case of mitonuclear discordance in the genus *Panthera* (Li et al., 2016), highlighting the need of using information from the nuclear genome to correctly infer the evolutionary relationships among species. Although the autosomal phylogenomic tree was based on complete genomes, low resolution within African and Asian clades was observed and monophyly was not recovered for some of the subspecies (Fig. 2A). Establishing the phylogenetic position of the Arabian leopard is critical for conservation programs, since even recently (Kitchener et al., 2017) the Arabian leopard was considered a consubspecific species together with the African *P. p. pardus*, as the mitochondrial inference indicates. Our genome-wide data advocates for the genomic distinction of the Arabian leopard, confirming that it needs to be managed as a separate conservation unit, as it has been done so far. Given that the Arabian leopard is the sister group to all other Asian subspecies, our findings suggest Arabia may have served as a stepping stone for the subsequent expansion across the rest of the Asian continent and perhaps Europe. Whole nuclear genomes of European individuals should be sequenced to resolve the complete evolutionary history of the species. All other phylogenomic relationships were found to be in line with previous phylogenomic results (Paijmans et al., 2021).

When estimating the past evolutionary history, a continuous trend of reduction in effective population size towards the present was observed for both Arabian samples (Fig. 3). Low heterozygosity levels for both Arabian leopards (Fig. 4A) and the medium coverage for both samples could influence the PSMC analysis, as this software relies on the heterozygosity levels to reconstruct the historical evolutionary history of each sample (Li & Durbin, 2011). Wild animals from different locations need to be sequenced at higher coverage to explore their evolutionary history and investigate the potential variability of population trends within the Arabian leopard populations.

As previously discussed by Pečnerová et al., (2021), African leopards present high levels of genetic diversity and high continent-wide genetic connectivity, considering its trophic position. Here, we replicated their findings showing that African leopards have the highest genome-wide heterozygosity levels and the lowest percentage of genome in RoH of all leopard populations (Fig. 4 and Table S2). Both Arabian leopards showed low heterozygosity levels (Table 1 and Fig. 4A). This heterozygosity could be consistent with a scenario of strong genetic drift acting upon the population, most likely due to the low long-term effective population size observed (Fig. 3) as well as inbreeding (Fig. 4). This might turn into a reduced ability to adapt to any rapid environmental change or the emergence of new diseases such as SARS-CoV-2 in leopards (Mahajan et al., 2022; Ralls et al., 2020). More than 50% of their genome is under RoH (Fig. 4B and Table S2), with different sizes of ROHs explaining long-term and recent inbreeding. Short and medium RoHs are features of populations that have experienced an old population bottleneck (Ceballos et al., 2018). Both Arabian leopards contained almost the same percentage of short and medium RoHs, indicating a similar evolutionary history (Fig. 4B and Table S2). These short and medium RoHs are consistent with the historical effective population reduction trends (Fig. 3), suggesting that this subspecies might have suffered a prolonged past bottleneck. Interestingly, the female Arabian2 had about 10% more recent and long RoHs than the male Arabian1 (Table S2). This may be explained by the captive breeding program, as long RoHs typically arise in consanguineous communities caused by recent inbreeding loops (Ceballos et al., 2018). This is an unexpected result, as in every breeding event inside the conservation program, there is planned inbreeding avoidance by mating with at least one wild-born individual. However, the founder population (wild-born individuals) could have been already related between them or coming from the same population, contributing to the increase in inbreeding. Only the highly inbred *P. p. orientalis* leopard from the Zoo shows similar levels of genome-wide heterozygosity and RoHs as the ones observed in the Arabian leopards. To avoid this situation any further, we strongly recommend incorporating genomic data to estimate relatedness between individuals in the ongoing leopard breeding program, as genomics, together with other disciplines, is an essential tool for the conservation success of target species. All other Asian samples were in line with previous results, with the Javan leopard (*P. p. melas*) from a small island population also showing low heterozygosity and high frequency of short RoHs and the Amur leopard with relatively low heterozygosity values and low percentage of the genome under RoH (Fig. 4, Fig. S5 and Table S2).

Despite the existence of morphological differences (Paijmans et al., 2021), we found introgression signals between Arabian and Anatolian (*P. p. tulliana*) leopards and between Arabian and African leopards (Fig. 5). Gene flow between big cats is a well-known phenomenon (Figueiró et al., 2017). Thus, the description of gene flow between geographically close subspecies of leopards is not surprising. On the one hand, the introgression between African and Arabian leopards is also partially supported by the mitochondrial phylogeny, as both groups cluster together, suggesting a possible migration corridor for females between Africa and the Arabian peninsula. However, we cannot rule out that this topology is due to mitogenomic incomplete lineage sorting, as in leopards males have higher dispersal than females (de Oliveira et al., 2022). On the other hand, the introgression between Arabian and Anatolian leopards is supported by the autosomal phylogeny, as the 100 Kbp sliding windows phylogeny clusters Arabian and one sample of Anatolian leopard together (Fig. S1B). Their past continuous distribution through Northern Arabia could have helped in promoting gene flow between these two genetically and morphologically distinct subspecies. Finally, we also found several other comparisons between subspecies of leopards to be significant, mainly between Asian subspecies, revealing a complex history of gene flow within the species (Table S4).

Genetic load is described as the resulting reduction in individual and mean population fitness due to deleterious mutations originated by mutation or gene flow and maintained or even increased by genetic drift or reduced efficacy of purifying selection (Bertorelle et al., 2022). In diploid organisms, genetic load can be separated (assuming recessivity) into the realised load (homozygous, expressed and with effects on the current generation) and the masked load (heterozygous, recessive deleterious mutations that can be expressed in future generations) (Bertorelle et al., 2022). Recently, several studies have reported purging of genetic load, especially in long-term isolated and inbred populations (Dussex et al., 2021; Kleinman-Ruiz et al., 2022). However, few studies on big cats have focused on reporting mutational load levels (but see de Manuel et al., 2020; Khan et al., 2021). In leopards, several genomic studies have recently focused on genomics and landscape ecology, but the incidence of mutational load across subspecies has not been evaluated yet (Paijmans et al., 2021; Pečnerová et al., 2021). Here, we found a reduced total genetic load (in the number of high-impact deleterious alleles) for both Arabian samples compared with African and the rest of Asian leopards (Fig. 6A). This result is also supported by the ratio of derived alleles (Fig. 6B). Interestingly, when high-impact, moderate-impact and low-impact deleterious alleles were split in homozygous or heterozygous derived alleles (as a proxy of realised and masked load), we observed a high realised load (with a large number of both alleles in the homozygous state) and a lower masked load (with less alleles in the heterozygous state) in Arabian leopards (Fig. 6A and Fig. S6), following the theoretical predictions after a prolonged bottleneck (Bertorelle et al., 2022). During population decline, the composition of genetic load changes, with many previously masked mutations becoming expressed and, as a consequence, increasing the realised load and decreasing the masked load (Bertorelle et al., 2022). We also observe an increase of the realised load and a decrease of the masked load for Asian (excluding Arabian) compared to African leopards (Fig. 6 and Fig. S6). However, conversely to Arabian leopards, the total load seems to have experienced a relaxation of the purging, possibly due to a posterior expansion. The Arabian leopard has been long-term isolated and together with the putative slow increase in inbreeding (and subsequent reduction of masked load) has possibly allowed the purging of high-impact deleterious alleles, mainly in the heterozygous state. As a consequence, around 40% and 60%, respectively, of the high-impact deleterious alleles were found within RoHs of both Arabian leopards (Fig. S7). During bottlenecks, inbreeding and drift increase homozygosity, turning masked load into realised load. Then, purifying selection acts upon the high-impact deleterious mutations but the moderate-impact deleterious mutations escape it and if the bottleneck is prolonged enough, some of them become fixed (Bertorelle et al., 2022). Interestingly, the Arabian leopard shows similar levels of moderate-impact and low-impact deleterious alleles compared to African and the rest of Asian leopards, as selection has not been as strong in these alleles. This is consistent with the purging of recessive high-impact deleterious alleles as a consequence of increased inbreeding (Bertorelle et al., 2022; Caballero et al., 2016). For instance, purging has been reported in some island populations of the Endangered Kakapo, with lower genetic diversity and population effective size and higher inbreeding and longer RoHs than other mainland populations (Dussex et al., 2021).

Inbreeding depression is defined as the increased homozygosity resulted from inbreeding, causing a reduction in fitness (Gooley et al., 2017). The genome-wide heterozygosity values for the two Arabian individuals were one of the lowest ever reported for the species and more than half of the genomes of the two Arabian individuals were within RoHs (Fig. 4). However, the amount of high-impact deleterious alleles was significantly lower and concordant with purging of deleterious mutations along a prolonged past bottleneck, as reported by evolutionary history analyses (Fig. 3 and Fig. 6), highlighting the importance of performing a comprehensive genomic study to evaluate the status of a species. If only heterozygosity values of the Arabian leopard were taken into account, the subspecies could be considered to be on the brink of an important inbreeding depression, but until now no reduction in fitness has been reported. Surprisingly, the out-of-RoH heterozygosity for both Arabian leopards is almost twice that of the genome-wide heterozygosity of the healthiest Asian individuals, highlighting that the areas without RoHs contain higher genetic diversity. Nonetheless, low genome-wide genetic diversity can result in health issues, causing a negative impact on reproductive fitness and life quality (Farquharson et al., 2018; Irizarry et al., 2016). In fact, genetic depletion has been already highlighted as one of the current threads for the Arabian leopard (Mallon & Budd, 2011). Overall, our results highlight that genomic tools are essential to assess the situation of endangered and elusive species. Genome-wide data successfully resolved the phylogenetic position of the Arabian leopard as sister to the rest of Asian subspecies, a topic highly discussed over the last two decades (Paijmans et al., 2021; Pečnerová et al., 2021; Uphyrkina et al., 2001). Moreover, using genomic data we provided accurate estimations of genetic diversity, population structure and demographic history for the Critically Endangered Arabian leopard, confirming a prolonged past bottleneck with subsequent inbreeding and purging of deleterious mutations. Ultimately, our study stresses the benefits of using genomic tools both from an evolutionary and a conservation perspective and highlights the importance of integrating the field of genomics when managing *in-situ* and *ex-situ* endangered and elusive species.

## Supporting information

Cross validation analyses for admixture from k=2 to k=5.

Percentage of short, medium, long and total RoHs along the genome per individual.

Number (NROH) and addition (SROH) of RoHs per individual.

Introgression analyses between all subspecies of leopards.

Phylogenomic consensus tree with different sizes of sliding windows.

PSMC analysis for the two Arabian leopards with 100 bootstrap.

PSMC of high-coverage downsampled individuals

Genome-wide out-of-RoH heterozygosity

Pearson correlation between the number (NROH) and the addition (SROH) of RoHs.

Moderate and low genetic mutational load in leopards.

## Acknowledgements

This work was supported by funding from the European Research Council (ERC) under the European Union’s Horizon 2020 research and innovation programme (grant agreement No. 864203), “Unidad de Excelencia María de Maeztu”, funded by the AEI (CEX2018-000792-M) to TMB, and grants PGC2018-098290-B-I00 (MCIU/AEI/FEDER, UE) Spain, PID2021-128901NB-I00 (MCIN/AEI/ 10.13039/501100011033 and by ERDF, A way of making Europe), Spain to SC. We acknowledge both Anders Albrechtsen and Nicolas Dussex for useful comments with introgression and mutational load analyses, respectively. We further acknowledge Johanna Paijmans for helping during the downloading of some leopard genomes. We are deeply thankful to Dr. Mohammed Qurban, chief executive officer of the National Center for Wildlife (NCW), and the former vice president of the formal Saudi Wildlife Authority Dr. Hani Tatwany for their support and encouragement since the beginning of the project. We are also grateful to all personnel involved in the Arabian Leopard program at the Royal Commission for AlUla and their partners in the Arabian Leopard project for support. GR was funded by an FPI grant from the Ministerio de Ciencia, Innovación y Universidades, Spain (PRE2019-088729), AT is supported by “la Caixa” doctoral fellowship program (LCF/BQ/DR20/11790007) and BB-C was funded by FPU grant from Ministerio de Ciencia, Innovación y Universidades, Spain (FPU18/04742).

## Data and code availability

The accession number and the code used for these analyses will be available after publication.

