## Supplementary material for "Genomics reveals introgression and purging of deleterious mutations in the Arabian leopard (*Panthera pardus nimr*)": Percentage of short, medium, long and total RoHs along the genome per individual.

**Table S2:** Percentage of short, medium, long and total RoHs along the genome per individual.

| Id | Short (<500 Kbp) | Medium (0.5-1 Mbp) | Long (>1 Mbp) | Total |
|---|---|---|---|---|
| PAR2 – Ghana | 3.7 | 3.5 | 5.5 | 12.7 |
| PAR4 – Namibia | 3 | 1.8 | 5 | 9.8 |
| PAR6 – TanzaniaN | 3.3 | 2.1 | 6.6 | 12 |
| NIM1 – Arab1 | 38.5 | 7.4 | 4.2 | 50.1 |
| NIM2 – Arab2 | 42.7 | 11.58 | 13.77 | 68.05 |
| FUS1 – Nepal | 7 | 3.7 | 5.6 | 16.3 |
| MEL – Java | 29.3 | 5.6 | 1.8 | 36.7 |
| ORI2 – Zoo | 18 | 12.5 | 20.6 | 51.1 |
| ORI3 – Amur | 12.8 | 4.9 | 4.3 | 22 |
