## Supplementary material for "Genomics reveals introgression and purging of deleterious mutations in the Arabian leopard (*Panthera pardus nimr*)": Number (NROH) and addition (SROH) of RoHs per individual.

**Table S3:** Number (NROH) and addition (SROH) of RoHs per individual.

| ID | Number of Rohs (NROH) | Addition of RoHs (SROH) |
|---|---|---|
| PAR2 – Ghana | 591 | 300,871,563 |
| PAR4 – Namibia | 466 | 232,193,737 |
| PAR6 – TanzaniaN | 524 | 284,141,682 |
| NIM1 – Arabian1 | 5,149 | 1,169,555,244 |
| NIM2 – Arabian2 | 5,442 | 1,586,284,559 |
| FUS1 – Nepal | 1,037 | 383,816,415 |
| MEL – Java | 3,790 | 858,606,362 |
| ORI2 – Zoo | 2,614 | 1,193,748,995 |
| ORI3 – Amur | 12.8 | 4.9 |
