## Supplementary material for "Genomics reveals introgression and purging of deleterious mutations in the Arabian leopard (*Panthera pardus nimr*)": Introgression analyses between all subspecies of leopards.

**Table S4:** Introgression analyses between all subspecies of leopards, following a tree-like model and using the lion as an outgroup. Z-scores higher than 3 are shown in bold.

| P1 | P2 | P3 | Outgroup | D-stat | Standard error | Z-score | BABA | ABBA | SNPs |
|---|---|---|---|---|---|---|---|---|---|
| *P. p. tulliana* | *P. p. nimr* | *P. p. pardus* | *P. leo* | -0.1122 | 0.002900 | **-38.682** | 196409 | 246046 | 16218841 |
| *P. p. fusca* | *P. p. nimr* | *P. p. pardus* | *P. leo* | -0.0645 | 0.003035 | **-21.256** | 235294 | 267746 | 16315748 |
| *P. p. melas* | *P. p. nimr* | *P. p. pardus* | *P. leo* | -0.0580 | 0.003247 | **-17.854** | 242200 | 272006 | 16290064 |
| *P. p. kotiya* | *P. p. nimr* | *P. p. pardus* | *P. leo* | -0.0572 | 0.003158 | **-18.098** | 235593 | 264158 | 16307189 |
| *P. p. orientalis* | *P. p. nimr* | *P. p. pardus* | *P. leo* | -0.0604 | 0.003159 | **-19.126** | 240721 | 271684 | 16315894 |
| *P. p. delacouri* | *P. p. nimr* | *P. p. pardus* | *P. leo* | -0.0663 | 0.003166 | **-20.932** | 240076 | 274152 | 16210368 |
| *P. p. fusca* | *P. p. tulliana* | *P. p. nimr* | *P. leo* | -0.2918 | 0.004855 | **-60.097** | 178604 | 325757 | 16218695 |
| *P. p. kotiya* | *P. p. tulliana* | *P. p. nimr* | *P. leo* | -0.2912 | 0.005172 | **-56.299** | 173971 | 316873 | 16210136 |
| *P. p. melas* | *P. p. tulliana* | *P. p. nimr* | *P. leo* | -0.3175 | 0.004915 | **-64.608** | 182818 | 352911 | 16193011 |
| *P. p. delacouri* | *P. p. tulliana* | *P. p. nimr* | *P. leo* | -0.3173 | 0.004910 | **-64.611** | 182300 | 351719 | 16113315 |
| *P. p. orientalis* | *P. p. tulliana* | *P. p. nimr* | *P. leo* | -0.3105 | 0.004852 | **-63.992** | 182992 | 347758 | 16218841 |
| *P. p. kotiya* | *P. p. fusca* | *P. p. tulliana* | *P. leo* | 0.0661 | 0.004214 | **15.697** | 177128 | 155147 | 16214345 |
| *P. p. melas* | *P. p. fusca* | *P. p. tulliana* | *P. leo* | -0.1957 | 0.004268 | **-45.854** | 142910 | 212469 | 16197220 |
| *P. p. delacouri* | *P. p. fusca* | *P. p. tulliana* | *P. leo* | -0.1727 | 0.004125 | **-41.857** | 145013 | 205540 | 16117524 |
| *P. p. orientalis* | *P. p. fusca* | *P. p. tulliana* | *P. leo* | -0.1603 | 0.003954 | **-40.545** | 144892 | 200224 | 16223050 |
| *P. p. melas* | *P. p. kotiya* | *P. p. fusca* | *P. leo* | -0.1570 | 0.005485 | **-28.627** | 189771 | 260473 | 16285634 |
| *P. p. delacouri* | *P. p. kotiya* | *P. p. fusca* | *P. leo* | -0.1046 | 0.005735 | **-18.233** | 202977 | 250383 | 16206034 |
| *P. p. orientalis* | *P. p. kotiya* | *P. p. fusca* | *P. leo* | -0.0875 | 0.005681 | **-15.394** | 206477 | 246057 | 16311398 |
| *P. p. delacouri* | *P. p. melas* | *P. p. kotiya* | *P. leo* | 0.0434 | 0.005509 | **7.871** | 139156 | 127592 | 16180653 |
| *P. p. orientalis* | *P. p. melas* | *P. p. kotiya* | *P. leo* | 0.0659 | 0.005125 | **12.858** | 142288 | 124696 | 16285780 |

| | | | | | | | | | |
|---|---|---|---|---|---|---|---|---|---|
| *P. p. orientalis* | *P. p. delacouri* | *P. p. melas* | *P. leo* | 0.0041 | 0.005376 | 0.769 | 140664 | 139507 | 16189130 |
