## Supplementary material for "Genomics reveals introgression and purging of deleterious mutations in the Arabian leopard (*Panthera pardus nimr*)": Phylogenomic consensus tree with different sizes of sliding windows.

Sliding windows of 500Kb

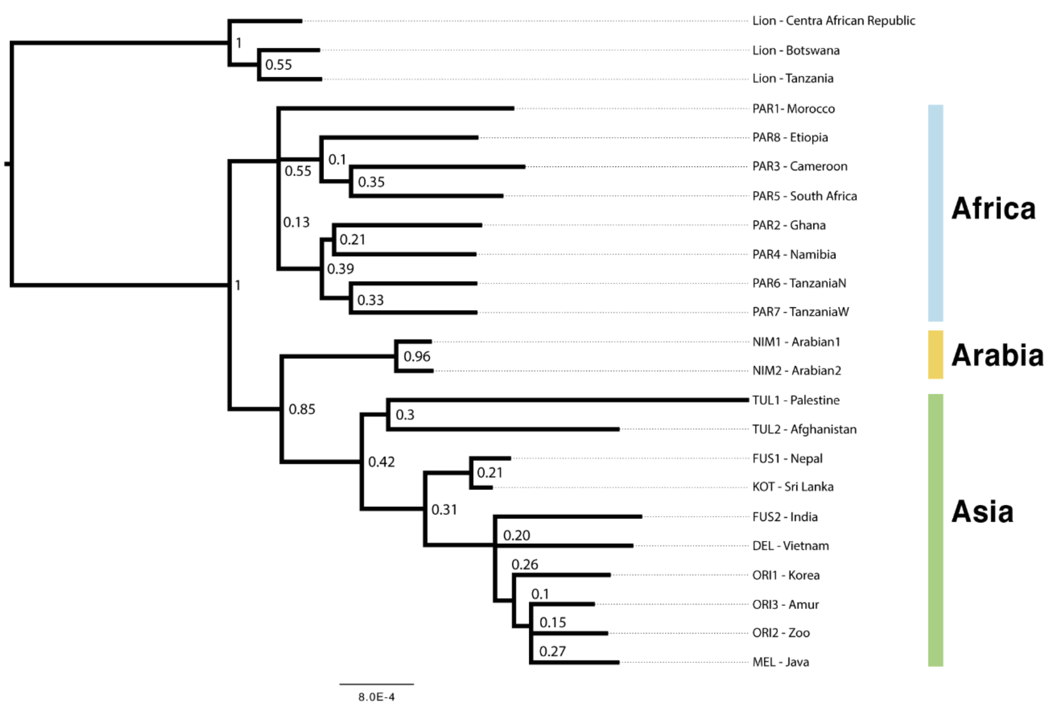


Sliding windows of 100Kb

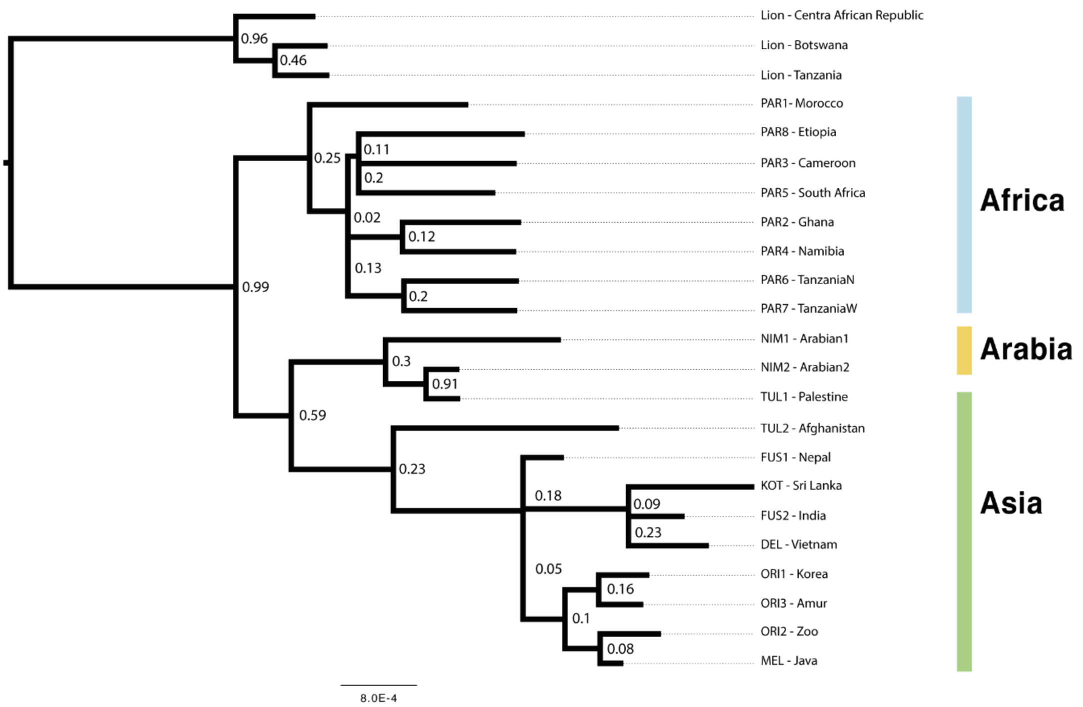


**Fig. S1:** Phylogenomic consensus tree with different sizes of sliding windows. (A) Phylogenomic consensus tree from 480 Maximum Likelihood trees of 500 Kbp non-overlapping sliding windows. (B) Phylogenomic consensus tree from 2,400 Maximum Likelihood trees of 100 Kbp non-overlapping sliding windows. In both cases, sliding windows were produced along the largest chromosome (240 Mbp), all currently accepted subspecies of leopards were present and the lion was used as an outgroup. Abbreviations are as follows: PAR, *P. p. pardus;* NIM, *P. p. nimr*; TUL, *P. p. tulliana*; FUS, *P. p. fusca*; KOT, *P. p. kotiya*; DEL, *P. p. delacouri*; ORI, *P. p. orientalis* and MEL, *P. p. melas*.
