## Supplementary material for "Genomics reveals introgression and purging of deleterious mutations in the Arabian leopard (*Panthera pardus nimr*)": PSMC analysis for the two Arabian leopards with 100 bootstrap.

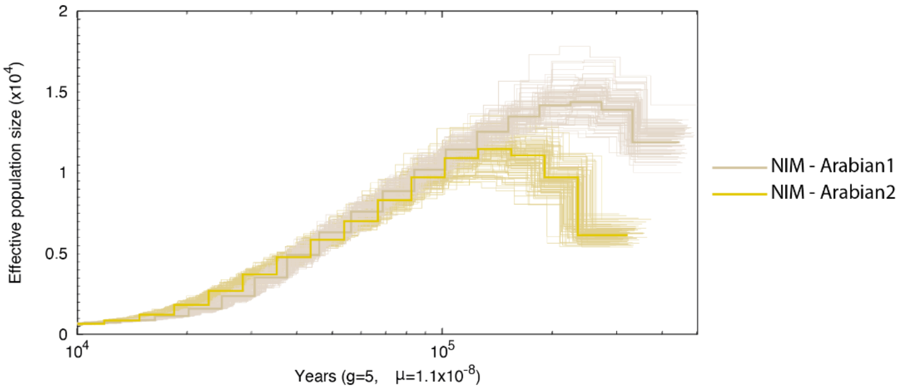


**Fig. S2:** PSMC analysis for the two Arabian leopards with 100 bootstrap for each sample. Generation time was set to 5 years and substitution rate to $1x10^{-8}$ per site per year. Abbreviation is as follows: NIM, *P. p. nimr*.
