## Supplementary material for "Genomics reveals introgression and purging of deleterious mutations in the Arabian leopard (*Panthera pardus nimr*)": PSMC of high-coverage downsampled individuals

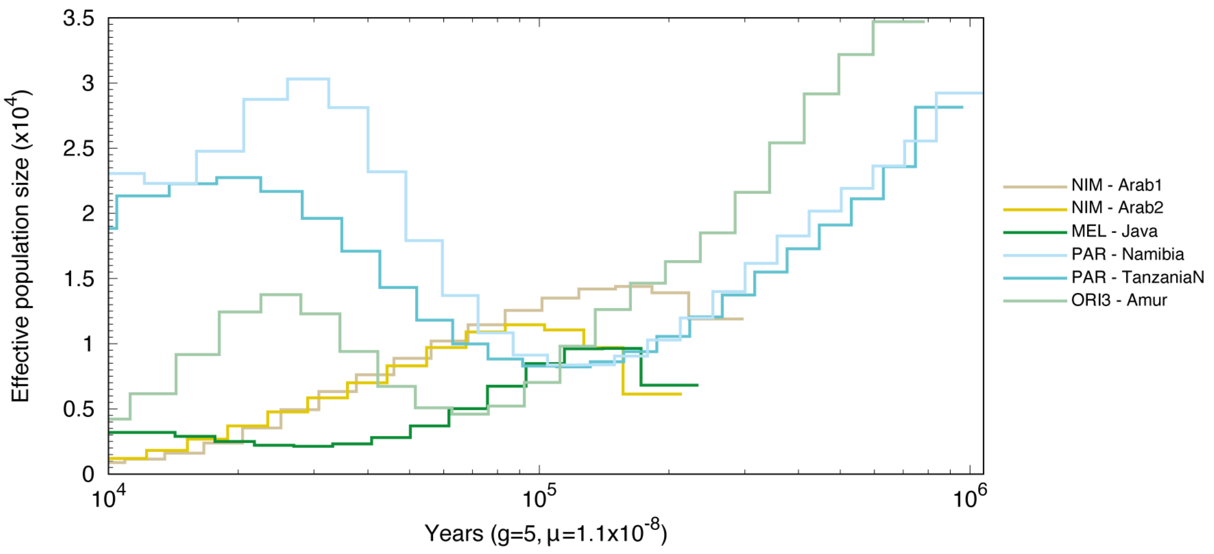


**Fig. S3:** PSMC of high-coverage individuals downsampled to similar coverage levels than the two Arabian leopards. Generation time was set to 5 years and substitution rate to $1x10^{-8}$ per site per year. Abbreviations are as follows: NIM, *P. p. nimr*; MEL, *P. p. melas*; PAR, *P. p. pardus* and ORI, *P. p. orientalis*.
