## Supplementary material for "Genomics reveals introgression and purging of deleterious mutations in the Arabian leopard (*Panthera pardus nimr*)": Genome-wide out-of-RoH heterozygosity

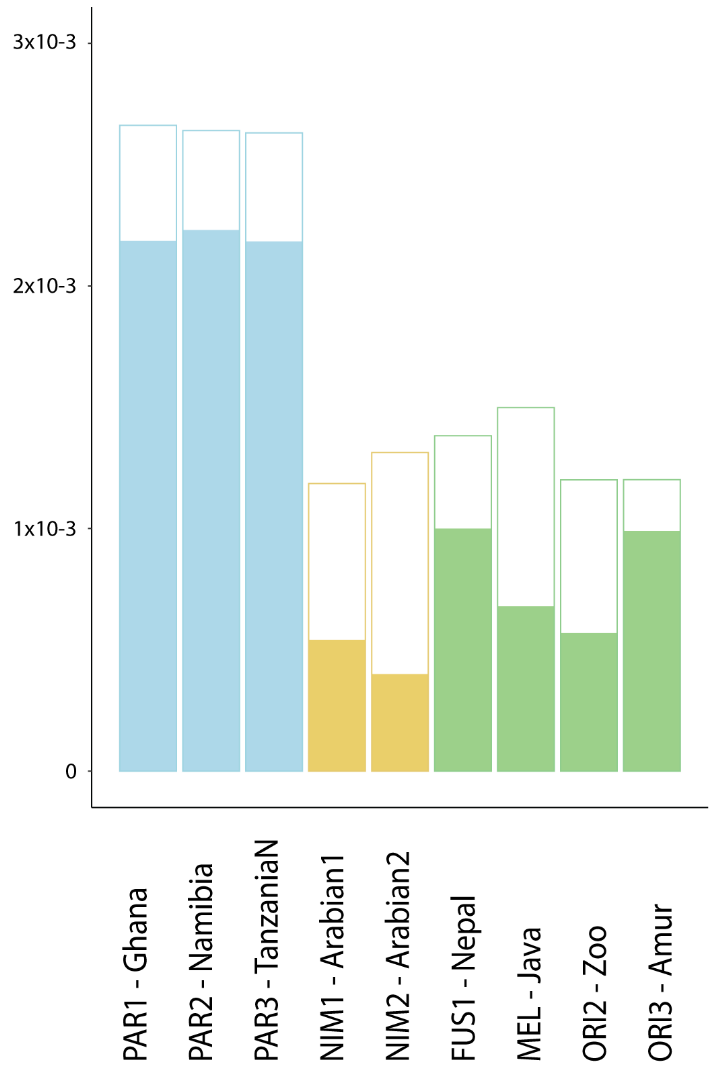


**Fig. S4:** Genome-wide heterozygosity levels for the high-coverage samples in light colours and genome-wide heterozygosity out-of-RoH in white, except for the sample from Nepal where out-of-RoH heterozygosity is lower than genome-wide heterozygosity. Abbreviations are as follows: PAR, *P. p. pardus;* NIM, *P. p. nimr*; FUS, *P. p. fusca*; MEL, *P. p. melas* and ORI, *P. p. orientalis*.
