## Supplementary material for "Genomics reveals introgression and purging of deleterious mutations in the Arabian leopard (*Panthera pardus nimr*)": Pearson correlation between the number (NROH) and the addition (SROH) of RoHs.

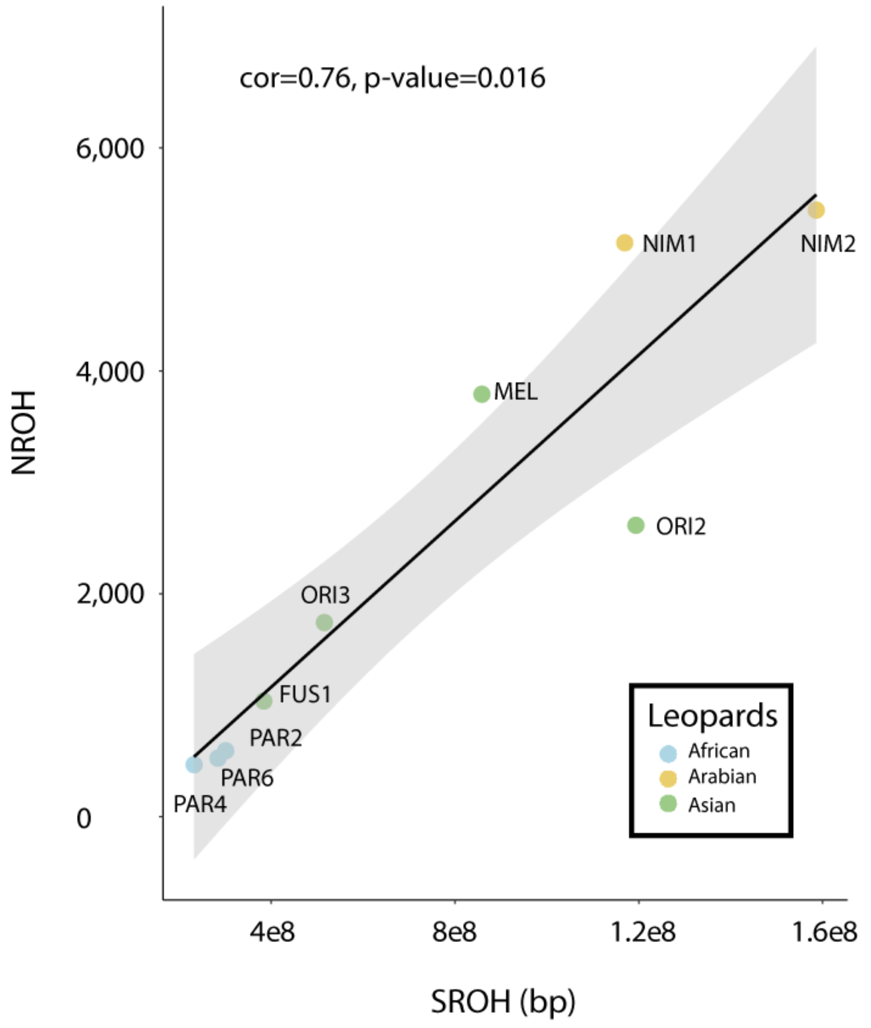


**Fig. S5:** Pearson's correlation between the number (NROH) and the addition (SROH) of RoHs. Line drawn only for representation reasons, as samples above the line will have experienced long-term inbreeding and samples below the line will have experienced recent inbreeding.
