## Supplementary material for "Genomics reveals introgression and purging of deleterious mutations in the Arabian leopard (*Panthera pardus nimr*)": Moderate and low genetic mutational load in leopards.

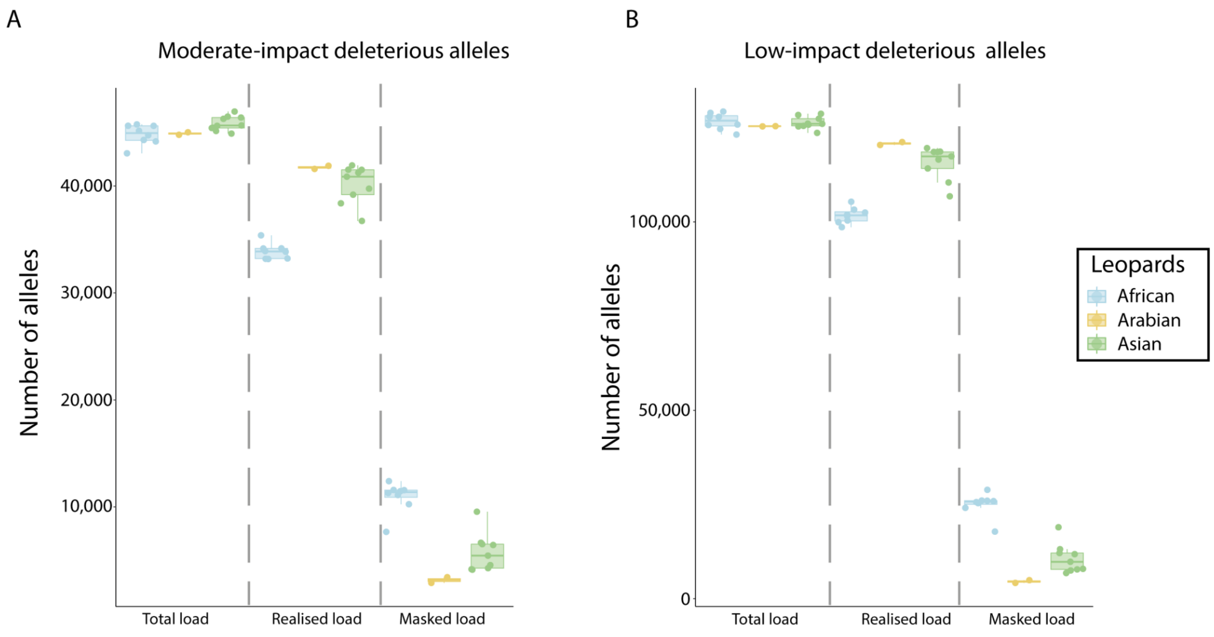


**Fig. S6:** Mutational load in leopards. (A) Number of moderate-impact deleterious alleles found in total, realised and masked genetic load for individuals from Africa, Arabia and the rest of Asia. (B) Number of low-impact deleterious alleles found in total, realised and masked genetic load for individuals from Africa, Arabia and the rest of Asia.
